# Functional analysis of KIT gene structural mutations causing porcine dominant white phenotype by using genome edited mouse models

**DOI:** 10.1101/688697

**Authors:** Guangjie Sun, Xinyu Liang, Ke Qin, Yufeng Qin, Xuan Shi, Peiqing Cong, Deling Mo, Xiaohong Liu, Yaosheng Chen, Zuyong He

**Affiliations:** State Key Laboratory of Biocontrol, School of Life Sciences, Sun Yat-Sen University, Guangzhou 510006, China

## Abstract

Dominant white phenotype in pigs is considered to be caused by two structural mutations in *KIT* gene, including a 450-kb duplication encompassing the entire *KIT* gene, and a splice mutation (G > A) at the first base in intron 17, which leads to the deletion of exon 17 in mature *KIT* mRNA, and the production of KIT protein lacking a critical catalytic domain of kinase. However, this speculation has not yet been validated by functional studies. Here, by using CRISPR/Cas9 technology, we created two mouse models mimicing the structural mutations of *KIT* gene in dominant white pigs, including the splice mutation mouse model *KIT ^D17/+^* with exon 17 of one allele of *KIT* gene deleted, and duplication mutation mouse model *KIT ^Dup/+^* with one allele of *KIT* gene coding sequence (CDS) duplicated. We found that each mutation individually can not cause dominant white phenotype. Splice mutation homozygote is lethal and heterozygous mice present piebald coat. Inconsistent with previous speculation, we found *KIT* gene duplication mutation did not confer the patched phenotype, and had no obvious impact on coat color. Interestingly, combination of these two mutations lead to dominant white phenotype. Further molecular analysis revealed that combination of these two structural mutations could inhibit the kinase activity of the KIT protein, thus reduce the phosphorylation level of PI3K and MAPK pathway associated proteins, which may be related to the observed impaired migration of melanoblasts during embryonic development, and eventually lead to dominant white phenotype. Our study provides a further insight into the underlying genetic mechanisms of porcine dominant white coat colour.

**Author summary:** KIT plays a critical role in control of coat colour in mammals. Two mutation coexistence in *KIT* are considered to be the cause of the *Dominant white* phenotype in pigs. One mutation is a 450-kb large duplication encompassing the entire *KIT* gene, another mutation is a splice mutation causing the skipping of KIT exon 17. The mechanism of these two mutations of KIT on coat color formation has not yet been validated. In this study, by using genome edited mouse models, we found each structural mutation individual does not lead dominant white phenotype, but combination of these two mutations could lead to a nearly complete white coat colour similar to pig dominant white phenotype, possibly due to the inhibition of the kinase activity of the KIT protein, thus its signalling function on PI3K and MAPK pathways, leading to impaired migration of melanoblasts during embryonic development, and eventually lead to dominant white phenotype. Our study provides a further insight into the underlying genetic mechanisms of porcine dominant white coat colour.

## Introduction

Due to domestication and long term selection, dominant white is a widespread coat color among domestic pig breeds, such as Landrace and Large White [1]. The dominant white phenotype in domestic pigs is considered to be caused by two structural mutations in the *KIT* gene, (1) a ∼450-kb tandem duplication that encompasses the entire *KIT* gene body and ∼150 kb upstream region of KIT gene and (2) a splice mutation at the first nucleotide of intron 17 in one of the *KIT* copies that leads to the skipping of exon 17, and the production of KIT protein lacking a critical region in kinase catalytic domain. [2–6].

KIT is a class III tyrosine kinase receptor, encoded by the *KIT* gene. KIT receptor is expressed on several cell types, including mast cells, hematopoietic progenitors, melanoblasts and differentiated melanocytes [7]. The binding of its ligand – stem cell factor (SCF) causes KIT to homodimerize, leading to the activation of its intrinsic kinase activity through autophosphorylation of tyrosine residues. KIT has a number of potential tyrosine phosphorylation sites, which interact with multiple downstream signaling pathways, including the PI3K, MAPK, and Src family kinase pathways [7, 8]. These pathways are involved in the regulation of cells growth, survival, migration and differentiation [9].

The 450-kb large duplication that encompasses the entire *KIT* gene body previously was speculated to confer the patch phenotype in pigs due to abnormal KIT expression [4]. Based on this, a hypothesis has been proposed that there is an evolutionary scenario whereby the duplication first occurred and resulted in a white-spotted phenotype that was selected by humans. The splice mutation occurred subsequently and resulted in a completely white phenotype, due to the skipping of exon 17 in the mature transcript removes a crucial part of the tyrosine kinase domain, thus enhances the defect in *KIT* signaling functions [5], and disturbs the migration of melanocyte precursors, leading to dominant white coat colour [2]. This seems reasonable, as normal migration and survival of neural crest-derived melanocyte precursors is dependent on KIT expression and the availability of its ligand [10]. Loss of function mutations in *KIT* gene could lead to white coat color in mouse, as documented in homozygous KIT ^K641E^ mouse [11] and KIT-deficient model W^v^/W^v^ [12]. However, functional analysis of the structural mutations in KIT gene of dominant pigs still need to be carried out to confirm the hypothesis. Here, by using CRISPR/Cas9 technology, we created mouse models mimicking the splice mutation and duplication mutation to investigate the underlying genetic mechanism of dominant white phenotype[13].

## Results

### Splice mutation but not the duplication mutation of *KIT* gene leads to altered coat color

In pigs, the wild-type KIT allele is recessive and denoted as *i*. Previous studies considered that two different mutant KIT alleles semidominant *I^P^* allele and the dominant *I* allele confer the patch and dominant white phenotype, respectively (Fig. 1A). *I* allele presents a 450-kb large duplication (two to three copies) that encompasses the entire *KIT* gene and at least one of the *KIT* copies carries a splice mutation (G>A at the first base in intron 17), causing exon skipping and the expression of a KIT protein lacking an essential part of the tyrosine kinase domain. *I^P^* allele involves the 450-kb duplication but not the splice mutation[6]. To investigate the effects of *KIT* gene structural mutations on coat color, we created three genome edited mouse models using C57BL/6 strain. The C57BL/6 mice is dominant black and broadly used in coat color study [14].

**Fig 1.**
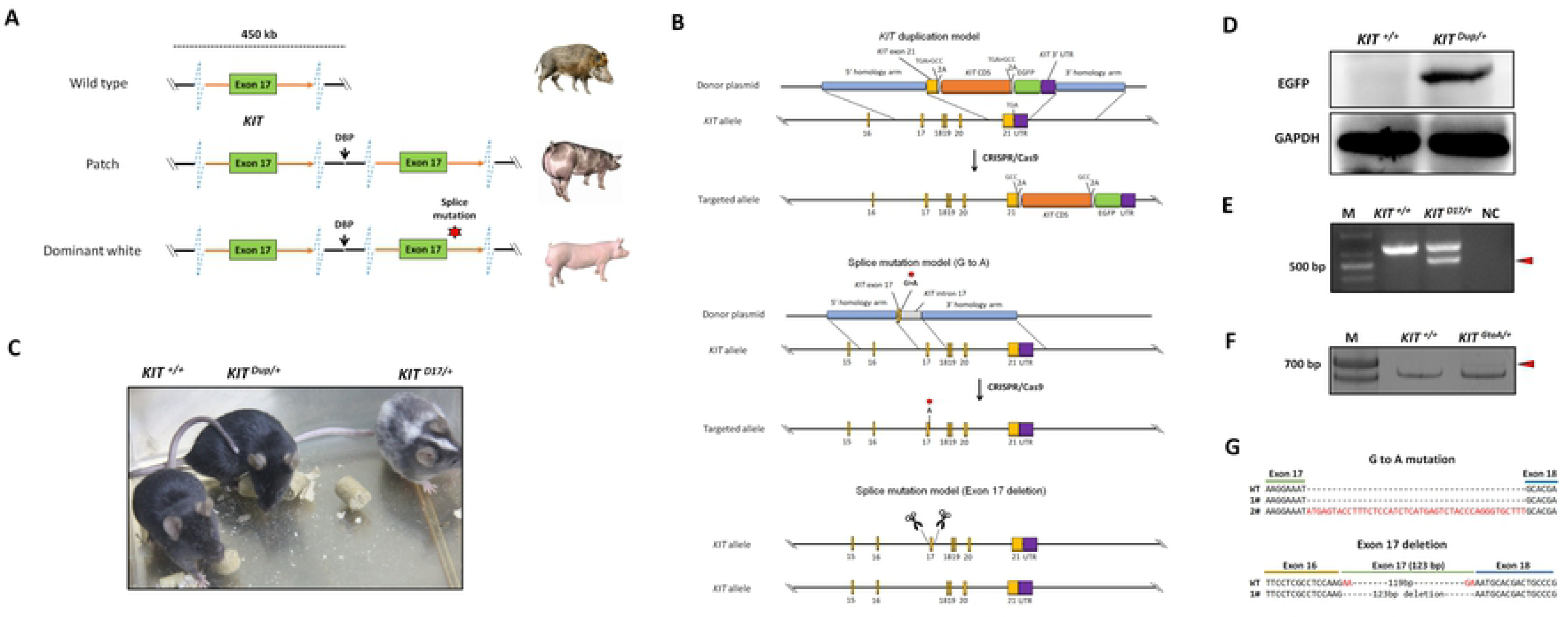
Generation of three mouse models mimicking structural mutations of KIT gene in dominant white pig. (A) The schematic summary of *KIT* mutations causing patch and dominant white phenotypes. of three coat phenotypes of pig. The patch coat colour is associated with a 450-kb large duplication that encompasses the entire KIT gene, and the dominant white is a associated with two to three copies KIT gene duplication and at least one of the KIT copies carries a splice mutation (G>A at the first base in intron 17), causing exon skipping and the expression of a KIT protein lacking an essential part of the tyrosine kinase domain. (B) To mimic the duplication mutation of KIT gene in patched pigs, the last exon of KIT gene with stop codon TGA mutated to GCC, linked with the CDS of KIT gene via a self-cleaving 2A peptide, followed by the linking with the enhanced green fluorescent protein (EGFP) reporter via 2A peptide was knocked in to the KIT locus through CRISPR/Cas9 mediated homologous recombination. To mimic the splice mutation, two mouse models were established. One has the first nucleotide (G) of KIT gene intron 17 substitute with A through CRISPR/Cas9 mediated homologous recombination, and another one with the exon 17 deleted by using paired sgRNA with one target the intron 16 and another one targets the intron 17. (C) Coat colour of the wild-type (*KIT^+/^*), heterologous of KIT duplication (*KIT ^Dup/+^*) and splice mutation (*KIT ^D17/+^*) mouse model. (D) Western blotting analysis confirmed the presence of EGFP expression in the skin of *KIT ^Dup/+^* mice, implying the inserted *KIT* CDS can be correctly expressed. (E) RT-PCR analysis of the deletion of exon 17 of *KIT* gene in *KIT ^D17/+^* mice. The truncated PCR product is indicated by arrow head. (F) RT-PCR analysis of the transcription product of *KIT* gene in *KIT ^GtoA/+^* mice. PCR product with insertion is indicated by arrow head. (G) Sanger sequencing of cDNA from F indicates exon 17 of *KIT* gene in *KIT ^GtoA/+^* mice is not removed, and a small percent of the transcript contained partial region of the intron 17. Sanger sequencing of cDNA from E indicates exon 17 is removed from the mature transcript in mature transcript in *KIT ^D17/+^* mice.

To mimic the duplication mutation (*I^P^* allele), we knocked in the CDS of KIT gene linked with the enhanced green fluorescent protein (EGFP) reporter via a self-cleaving 2A peptide to facilitate subsequent identification (Fig. 1B). The heterologous of *KIT* duplication mouse model was denoted as *KIT ^Dup/+^*. Western blot analysis results demonstrated that EGFP was extensively expressed in the skin of *KIT ^Dup/+^* mice, as compared with the wild-type control *KIT^+/+^*, which implying the inserted *KIT* CDS was correctly expressed, as 2A peptide strategy allows the co-expression of KIT proteins and EGFP from the integrated single vector (Fig. 1D). We found duplication of *KIT* gene did not result in the patch phenotype, as no obvious difference was observed on coat color between *KIT ^Dup/+^* and *KIT^+/+^* mice (Fig. 1C).

To mimic the splice mutation, we substitute the first nucleotide (G) of *KIT* gene intron 17 with A through CRISPR/Cas9 mediated homologous recombination (Fig. 1B). The heterologous of this splice mutation mouse model was denoted as *KIT ^GtoA/+^*. An extensive screening of the offspring implies that homologous of splice mutation of KIT gene could be lethal, as no survived individual of *KIT ^GtoA/GtoA^* has been identified. Big white spots appeared on the abdomen of *KIT ^GtoA/+^* mice as compared with *KIT^+/+^* (S1 Fig). To determine whether the G to A mutation at the first nucleotide of intron 17 of *KIT* gene can lead to the skipping of exon 17, RT-PCR was carried out by using the RNA isolated from the skin of *KIT ^GtoA/+^* mice. To our surprise, the results showed that exon 17 was not removed from the transcript mRNA, and a small percent of the transcript contained partial region of the intron 17, as determined by Sanger sequencing (Fig. 1F & G). As this model does not mimic the splice mutation well, we did not use it in the subsequent studies.

Therefore, in order to create a mouse model mimicking the skipping of exon 17, we have directly deleted the exon 17 in genomic by using paired sgRNA with one target the intron 16 and another one targets the intron 17 (Fig. 1B). The heterologous of this splice mutation mouse model was denoted as *KIT ^D17/+^*. An extensive screening of the offspring implies that homologous of splice mutation of *KIT* gene could be lethal, as no survived individual of *KIT ^D17/D17^* has been identified. In addition, IVF experiment by fertilization oocytes from *KIT ^D17/+^* females with sperm from *KIT ^D17/+^* males resulted in no survived individual of *KIT ^D17/D17^* can be obtained (S1 Table). This confirms the previous speculation that *I^L^* allele (single copy of *KIT* gene with the splice mutation) could be lethal in pigs, as this allele was not found among worldwide pig population [6]. RT-PCR analysis of the skin tissue from *KIT ^D17/+^* mice indicates that exon 17 is removed from the mature transcript (Fig. 1E), and this was confirmed by Sanger sequencing (Fig. 1G). Interestingly, compared with *KIT^+/+^*, *KIT ^D17/+^* mice presented a piebald coat colour in head and trunk, a vertical white stripe on the forehead, a half loop of white hair on the shoulder blade area, and dominant white at entire abdominal part. (Fig. 1D & S1 Fig).

### Splice mutation but not the duplication mutation of *KIT* gene significantly reduces melanin accumulation

Histological analysis (Fontana-Masson staining) of the back skin of 5-week old mice, revealed that similar to the *KIT ^+/+^* control mice, the hair follicles of both *KIT ^Dup/+^*and *KIT ^D17/+^* mice are long in length, and in the hair bulb, the dermal papilla is completely coated by matrix, the keratogenous zone region is clearly visible and is connected to hair shaft and dermal papilla, and fibrous tract is substantially invisible in the skin (Fig. 2). These results indicate that similar to the *KIT ^+/+^* control mice, the hair follicles of 5-week-old *KIT ^Dup/+^*and *KIT ^D17/+^* mice are in the growing stage (anagen V or VI), which is advantageous for the observation of the hair follicle shape, and melanin distribution due to melanin synthesis is more active during this stage [15]. The results of hair follicle shape imply that both the splice mutation and duplication mutation did not impair the hair follicle development significantly.

**Fig 2.**
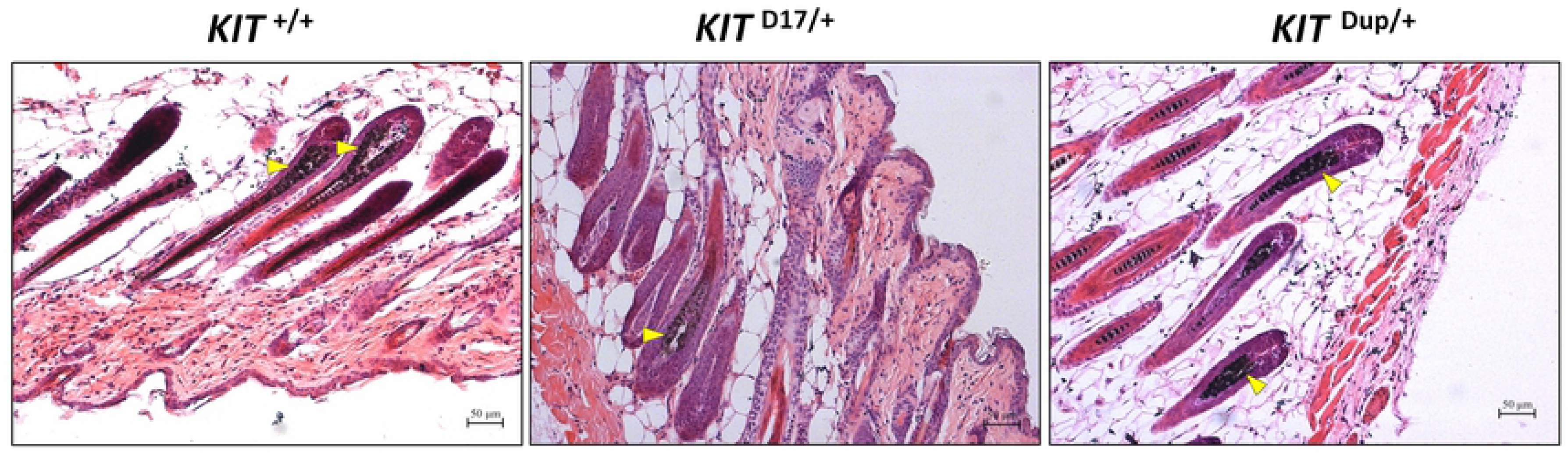
Histological analysis of the back skin of 5-week old *KIT ^+/+^, KIT ^D17/+^* and *KIT ^Dup/+^* mice. Melanin is stained with Fontana-Masson, and indicated by arrow head. Scale bar = 50 μM.

No obvious difference was observed in the content of melanin contained in hair follicles between *KIT ^Dup/+^* and *KIT ^+/+^* mice (Fig. 2). While almost no melanin is observed in the hair follicles within the white coat area, and the melanin level of hair follicles in the black coat area of *KIT ^D17/+^* is significantly lower (indicated by yellow arrow head). Therefore, the *KIT* duplication mutation did not impair the melanin accumulation, whereas the splice mutation significantly impaired melanin accumulation in hair follicle.

### The piebald coat colour of *KIT ^D17/+^* mice is caused by the reduction of melanocytes

The reduction of melanin content in hair follicles may be due to reduced melanocytes, or reduced ability of melanocytes on melanin synthesis. To determine the piebald coat of *KIT ^D17/+^* mice is caused by which factor, we used KIT protein as the marker to detect the distribution and amounts of melanocytes in the hair follicle of 5-week-old mice. Compared with the *KIT ^+/+^* control mice, we found that the immunohistochemical staining of KIT decreased in the hair follicles of black coat area of *KIT ^D17/+^* mice, while the staining decreased more significantly in the white coat region (Fig. 3A). This was confirmed by qPCR and Western blot analysis of the skin tissues (Fig. 3B and 3C). In theory, the deletion of exon 17 should not affect the expression level of *KIT* gene, thus the reduced *KIT* expression level in skin tissue should be due to reduced melanocyte quantity. qPCR analysis of the isolated peritoneal cell derived mast cells indicated that deletion of the exon 17 did not affect the expression level of *KIT* gene (Fig. 3B & S2 Fig). The reduced expression of *KIT* gene together with another two marker genes (DCT and Melan A) of melanocytes in *KIT ^D17/+^* mice (Fig. 3C) suggested that the splice mutation of *KIT* gene could lead to reduced number of melanocytes in mice hair follicles.

**Fig 3.**
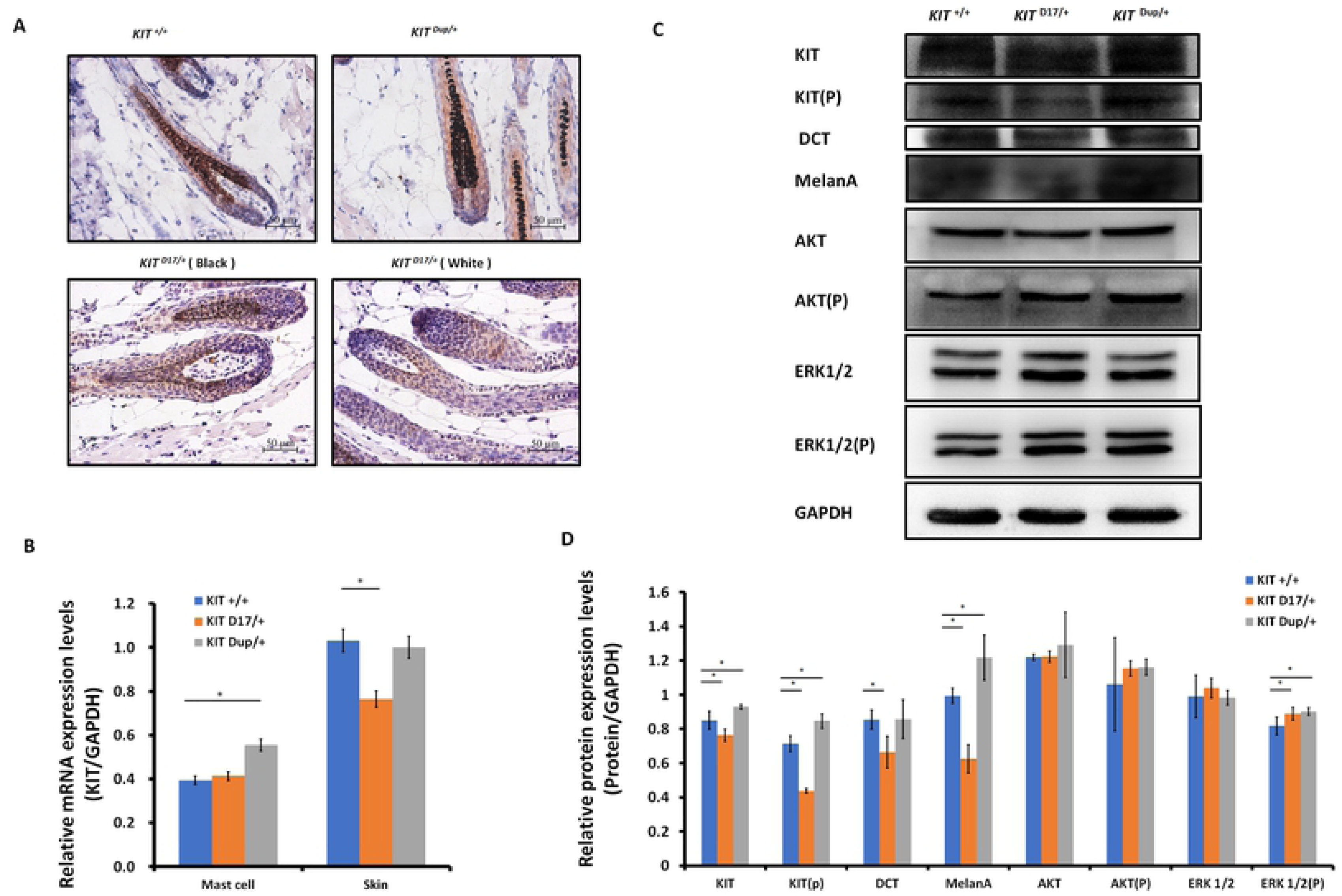
KIT splice mutation causes the reduction of melanocytes. (A) Expression of KIT protein in hair follicle of wild-type and mutant mice was determined by immunohistochemical staining. Melanin is stained with Fontana-Masson. Scale bar = 50 μM. (B) Transcriptional level of *KIT* gene in mast cell and skin tissue of wild-type and mutant mice was determined by qPCR analysis. (C) Expression level of KIT, DCT, MelanA, AKT, ERK1/2 and phosphorylation level of KIT, AKT, ERK1/2 in skin tissue of wild-type and mutant mice was determined by Western blot analysis. (D)Statistical analysis of relative protein expression levels base on the intensity of bands in C. * stands for p < 0.05.

Unlike the *KIT ^D17/+^*mice, the *KIT ^Dup/+^* mice contained an additional copy of *KIT* CDS, which in theory could lead to increased expression of *KIT* gene. qPCR analysis of the isolated mast cells confirmed that the expression level of *KIT* gene was improved in *KIT ^Dup/+^* mice as compared with the *KIT ^+/+^* control mice (Fig. 3B). However, in mouse skin, the expression level of KIT in *KIT^Dup/+^* mice was not significantly different from the *KIT ^+/+^* control mice as revealed by immunohistochemical and Western blot analysis (Fig. 3A & C). in addition, the expression level of melanocyte marker gene DCT was not affected, but another melanocyte marker gene MelanA was significantly improved in the skin of *KIT ^Dup/+^* mice (Fig. 3C). These results suggested that the duplication mutation of *KIT* gene may not affect the number of melanocytes in the skin, but may affect melanin synthesis.

Interestingly, we observed that the distribution of melanocytes in hair bulb is broader in *KIT ^D17/+^*mice as compared with the *KIT ^+/+^* control mice. This phenomenon was more obvious in white coat area than in the black coat area (Fig. 3A). In addition, we found the distribution of melanocytes in the hair bulb of *KIT ^Dup/+^* mice was relatively broader than that in the *KIT ^+/+^* control mice (Fig. 3A). Previous studies considered that only melanocytes that are close to the dermal papilla can secrete and provide melanin to the hair [16], we speculate the altered distribution of melanocyte in both *KIT* splice mutation and duplication mutation mice may have certain impact on melanin accumulation.

### Splice mutation of *KIT* gene impairs the kinase activity of the KIT protein and affects embryonic melanoblast migration

Previous studies speculated that *KIT* mutations in dominant white pigs could disturb the migration of melanocyte precursors melanoblasts during the embryonic period [2]. In order to determine whether the splice mutation or the duplication mutation of *KIT* gene impairs the migration of melanoblasts during embryonic period, we stained the *KIT* protein as a marker to detect the distribution of melanoblasts in the transverse section mice at E14.5. We found no obvious changes in the location of melanoblasts in *KIT ^Dup/+^*compared to the *KIT ^+/+^* control mice (S3 Fig). Though the distribution of melanoblasts in *KIT ^D17/+^* near the neural tube is not different between in *KIT ^D17/+^* and the *KIT ^+/+^* mice, the number of melanoblasts in the dorsolateral migration pathway, near the forelimb and the abdomen epidermis is significantly reduced (Fig. 4A). This indicates that duplication mutation of *KIT* gene alone does not impair the migration of melanoblasts, in contrast, the splice mutation could significantly impair the migration of melanoblasts at embryonic stage, however, it does not completely block the migration process, a certain number of melanoblasts could migrate to the corresponding destination positions, and leads to the piebald phenotype.

**Fig 4.**
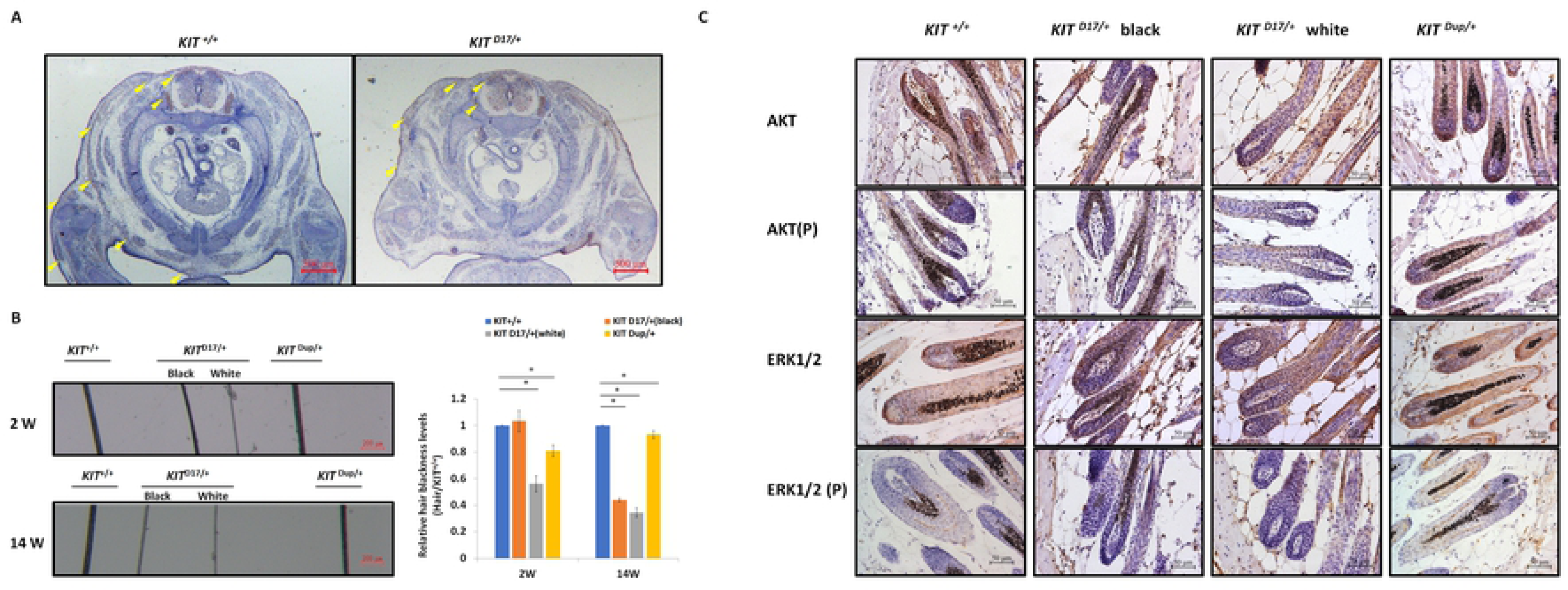
*KIT* spice mutation affects embryonic melanoblast migration and melanin accumulation. (A) KIT is used as marker to detect melanoblast migration in *KIT ^+/+^* and *KIT ^D17/+^* mice embryo (E14.5). Migrating melanoblasts were indicated by arrow heads. Scale bar = 500 μM. (B) Observation of hair from 2-W and 14-W old wild-type and mutant mice under a stereo microscope (left panel). Scale bar = 200 μM. The relative blackness of hair was quantified based on the intensity of image (right panel). * stands for p < 0.05. (C) AKT and ERK 1/2 expression and phosphorylation level in hair follicle of wild-type and mutant mice is determined by immunochemical staining. Scale bar = 50 μM.

We observed that, compared with the *KIT ^+/+^* control mice, the colour of black hair of *KIT ^D17/+^* mice became significantly lighter as the mice grew older. The determination of the blackness of mouse hair showed that the blackness of the black hair of *KIT ^D17/+^* mice was comparable to that of *KIT ^+/+^*mice at 2 W, however, the blackness of the black hair of *KIT ^D17/+^* mice decreased dramatically at 14 W, and was close to that of white hair of the *KIT ^D17/+^* mice (Fig. 4B). Interestingly, we found the blackness of the hair of *KIT ^Dup/+^*mice was relatively lower than that of *KIT ^+/+^*mice at 2 W, but became comparable to that of *KIT ^+/+^*mice at 14 W (Fig. 4B). Thus, we speculate that the splice mutation of *KIT* gene may affect the renewal or melanin synthesis function of melanocytes in mice, which in turn causes the blackness of hair to decrease rapidly with age. To examine whether the impaired melanoblast migration and melanocyte renewal is caused by altered kinase activity of the KIT protein receptor, Western blot analysis of the skin tissue was carried out. We found splice mutation significantly reduced the phosphorylation level of KIT protein (Fig. 3C & D), indicating this mutation could lead to impaired autophorylation ability of KIT protein. However, both the expression level and phosphorylation level of AKT, a key protein of the PI3K pathway, was not impaired by the splice mutation of *KIT* gene. Also, the expression of ERK1/2, key proteins of the MAPK pathway, was not affected by the splice mutation, but the phosphorylation level of ERK1/2 was slightly increased (Fig. 3C & D). This result looks confusing, therefore, we further analyzed the expression and phosphorylation levels of AKT and ERK1/2 in follicles by immunohistochemical (IHC) analysis. The amount of target protein was determined by using Combined Positive Score (CPS), which is the number of target protein staining cells divided by the total number of viable cells, multiplied by 100. The results revealed that the expression level of both AKT and ERK1/2 in hair follicles was not affected by the splice mutation of KIT gene (Fig. 4C), however, the phosphorylation levels of these proteins decreased significantly in the follicle of black coat region of *KIT ^D17/+^*mice, and phosphorylated AKT and ERK1/2 barely can be detected in the follicle of white coat region of *KIT ^D17/+^*mice (Fig. 4C). Western blot results indicate that the duplication mutation of *KIT* gene increased the phosphorylation level of KIT protein (Fig. 3C & D). This is probably due to the increased expression level of KIT protein. Similar to the splice mutation, the duplication mutation did not affect both the expression level and phosphorylation level of AKT. It also did not affect the expression level of ERK1/2, but slightly increased the phosphorylation level of ERK1/2 (Fig. 3C & D). IHC analysis revealed that both the expression level and the phosphorylation levels of AKT and ERK1/2 in the follicle of *KIT ^Dup/+^*mice was not significantly affected (Fig. 4C). As AKT and ERK1/2 are respectively involved in the PI3K and MAPK pathways, which are responsible for melanoblast migration and differentiation, and melanin synthesis in melanocyte, therefore, the impaired melanoblast migration and accelerated hair greying in *KIT ^D17/+^*mice should be related to impaired function of KIT kinase caused by the splice mutation of *KIT* gene.

### Combination of the splice mutation and duplication mutation of *KIT* gene caused severely impaired melanoblast migration during embryonic stage, and dominant white phenotype

As the splice mutation and duplication mutation of *KIT* gene individually did not lead to dominant white phenotype, we are curious whether the combination of these two mutations (denoted as compound mutations) can lead to dominant white phenotype. Therefore, the *KIT ^Dup/+^*male and the *KIT ^D17/+^*female was crossed to produce the *KIT ^Dup/D17^* offspring as determined by PCR analysis of the deleted exon 17 and integrated EGFP reporter (Fig. 5A). Interestingly, the *KIT ^Dup/D17^* mice presented a coat colour resembling the porcine dominant white phenotype: except for few gray hairs appearing near the eyelids and hip, the whole body was covered with white hairs (Fig. 5B). With the increase of age, the gray hairs of the eyelids and hips of *KIT ^Dup/D17^* mice gradually disappeared (S4 Fig). Through histological observation of the back skin of *KIT ^Dup/D17^* mice, we found that melanin is hardly visible in the hair follicles (Fig. 5C). However, no apparent morphological difference of the follicle was observed between *KIT ^Dup/D17^* mice and the *KIT ^+/+^* control mice. The hair follicles of 5-week-old *KIT ^Dup/D17^*mice showed a typical characteristics of the growing stage (anagen V or VI) (Fig. 5C). This indicates that the compound mutation did not affect the development of hair follicle, but severely impaired the accumulation of melanin in hair follicles.

**Fig 5.**
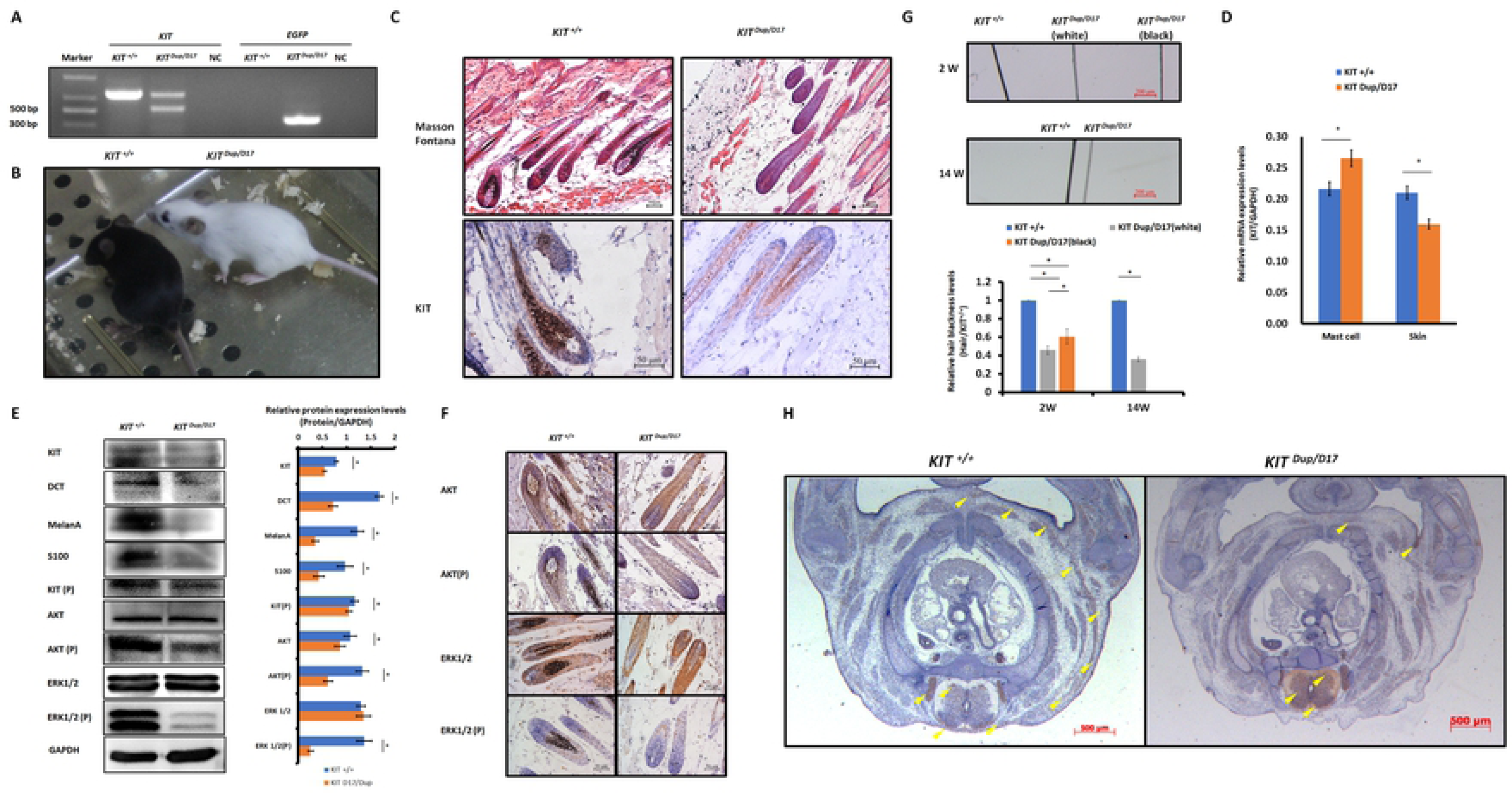
Combination of *KIT* duplication and splice mutation causes severely impaired KIT signaling function and melanoblast migration in embryonic stage. (A) Identification of *KIT ^Dup/D17^* mice through PCR analysis. (B) Coat colour of *KIT^+/+^* and *KIT ^Dup/D17^* mouse. (C) Histological analysis of melanin accumulation in hair follicle of *KIT^+/+^* and *KIT ^Dup/D17^* mice by Masson Fontana staining, and presence of melanocytes in hair follicle by immuostainging of KIT. Scale bar = 50 μM. (D) KIT mRNA levels mast cell and skin of *KIT^+/+^* and *KIT ^Dup/D17^* mice were determined by qPCR analysis. (E) Expression levels of KIT, DCT, MelanA, S100, AKT, ERK1/2 and phosphorylation levels of KIT, AKT, ERK1/2 in skin of *KIT^+/+^* and *KIT ^Dup/D17^* mice skin were determined by Western blot analysis (left panel) and quantified base intensity of bands (right panel). * stands for p < 0.05. (F) Expression and phosphorylation level of AKT and ERK 1/2 in hair follicle of *KIT^+/+^* and *KIT ^Dup/D17^* mice was determined by immunohistochemical analysis. Scale bar = 50 μM. (G) Observation of hair from 2-W and 14-W old of *KIT^+/+^* and *KIT ^Dup/D17^* mice under a stereo microscope (upper panel). Scale bar = 200 μM. The relative blackness of hair was quantified based on the intensity of image (lower panel). * stands for p < 0.05. (H) KIT is used as marker to detect melanoblast migration in *KIT ^+/+^* and *KIT ^Dup/D17^* mice embryo (E14.5). Migrating melanoblasts were indicated by arrow heads. Scale bar = 500 μM.

We suspected that similar to *KIT ^D17/+^*mice, the decreased melanin accumulation in the hair follicles of *KIT ^Dup/D17^* mice may be caused by reduced number of melanocytes in the hair follicles. qPCR analysis of the isolated mast cells from *KIT ^Dup/D17^* mice showed that compound mutations lead to improved expression level of *KIT* gene (Fig. 5D), which was mainly due to integrated additional copy of *KIT* CDS. In contrast, the expression level of *KIT* gene decreased significantly in skin tissue of *KIT ^Dup/D17^* mice as determined by qPCR analysis (Fig. 5D), this was further confirmed by Western blot analysis (Fig. 5E). The reduced expression of *KIT* gene together with another three marker genes (DCT, MelanA and S100) of melanocyte in skin tissue of *KIT ^Dup/D17^* mice (Fig. 5E) suggested that compound mutations of *KIT* gene could lead to reduced number of melanocytes in mice hair follicles, which may contribute to the decreased melanin accumulation in the hair follicles of *KIT ^Dup/D17^* mice.

IHC analysis of the skin tissue confirmed that the number of melanocytes in the hair follicles of *KIT ^Dup/D17^* mice decreased significantly as compared with the *KIT ^+/+^* control mice, with only a few layers of melanocytes in close proximity to the dermal papilla visible (Fig. 5C).

Few gray hairs presented in the whole white background of *KIT ^Dup/D17^* mice at young age, and they disappear gradually as determined by the hair blackness analysis (Fig. 5G & S4 Fig). This implies that the compound mutations of *KIT* gene may affect the renewal of melanocytes in hair follicles.

To investigate the underlying molecular mechanism of the compound mutations of *KIT* gene on coat colour changing, the kinase function of KIT protein was determined by Western blot analysis of the skin tissue of *KIT ^Dup/D17^* mice. The results showed that the phosphorylation level of KIT in the skin of *KIT ^Dup/D17^* mice is significantly lower than that of the *KIT ^+/+^* control mice (Fig. 5E). Both the expression level and phosphorylation level of AKT, a key protein of the PI3K pathway, decreased significantly. Though the expression of ERK1/2, key proteins of the MAPK pathway, was not affected, but the phosphorylation level of ERK1/2 decreased dramatically (Fig. 5E). These results indicate that the compound mutations of *KIT* gene substantially impaired the signaling function of KIT protein receptor on PI3K and MAPK pathways. However, to our surprise, further IHC analysis of the hair follicle showed that phosphorylation of AKT and ERK1/2 in *KIT ^Dup/D17^*seems not affected by the compound mutations as determined by CPS (Fig. 5F). This may imply a complex interaction between the splice mutation and duplication mutation of the *KIT* gene.

The compound mutations of *KIT* gene impaired the signaling function of KIT protein, which in turn could affect melanoblast migration at embryonic stage. Therefore, IHC analysis of the distribution of melanoblasts in the transverse section of *KIT ^Dup/D17^* mice at E14.5 by staining the marker protein KIT. The results showed that compared with the *KIT ^+/+^*control mice, the number of melanblast in the embryo of *KIT ^Dup/D17^* mice increased significantly in the neural tube, and the number of melanoblasts in the dorsolateral migration pathway, the forelimb epidermis and the abdomen epidermis decreased significantly (Fig. 5H). This result indicates that the compound mutations of *KIT* gene severely blocked melanoblast migration at embryonic stage, leading to increased accumulation of melanoblasts in the neural tube of the *KIT ^Dup/D17^* mice, and a coat colour resembling porcine dominant white phenotype.

## Discussion

In our study, we created mice models by using CRISPR/Cas9 technology to mimic the structural mutations of *KIT* gene in dominant white pigs. We used *KIT ^D17/+^* mice model to research the effect of coat colour on *KIT* exon 17 deletion and explored the impact mechanism of *KIT* duplication on coat colour by *KIT ^Dup/+^* mice model. We found that the *KIT* duplication did not influence mouse coat colour but *KIT* exon 17 deletion turns black hair of mouse into piebald colour. The experimental results prove that the *KIT* exon 17 deletion reduced the kinase function of KIT and impair it signaling transduction on PI3K and MAPK pathways, which are involved in melanoblast migration, leading to certain percent of melanoblast blocked in migration from dorsal to ventral region during embryo development, resulting in a piebald coat of the mouse. Interestingly, combination of these two mutations lead to dominant white phenotype. In mutation *KIT ^Dup/D17^* mouse embryo, melanoblasts severe blocked in the neural tube that could not migration. Those make *KIT ^Dup/D17^* mouse displaying *domiant white* phenotype (Fig 6).

**Fig 6.**
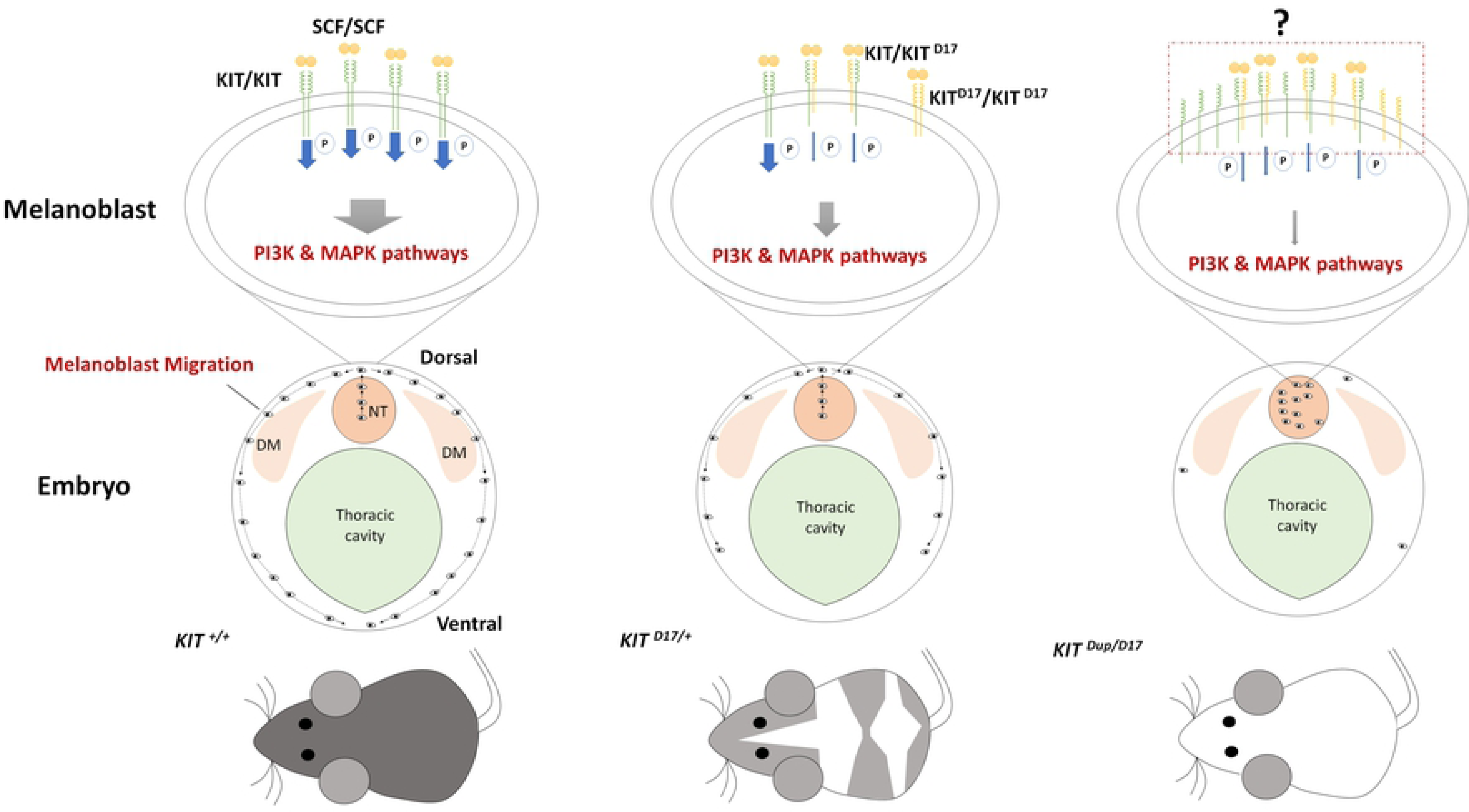
Schematic summary of the mechanism of KIT structural mutations on coat colour changing. SCF ligand may induce the formation of KIT/KIT ^D17^ dimer in melanoblast of mouse with KIT splice mutation (*KIT ^D17/+^* mouse), and reduced the kinase function of KIT and impair its signaling transduction on PI3K and MAPK pathways, which are involved in melanoblast migration, leading to certain percent of melanoblast blocked in migration from dorsal to ventral region during embryo development, resulting in a piebald coat of the mouse. Improved expression of normal form of KIT protein in mouse with the combination of KIT splice mutation and duplication mutation (*KIT ^Dup/D17^* mouse) may increase the chance of formation of KIT/KIT ^D17^ dimer upon the binding of SCF as compared with that in *KIT ^D17/+^* mice. Given that the amount of SCF ligand is limited, more KIT/KIT ^D17^ dimer presented on the cell surface of melanoblast may significantly reduce it signaling functions, resulting in more severely impaired melanoblast migration, with most melanoblast remaining in the neural tube, and resulting in a completely white coat colour. DM: dermamyotome; NT: neural tube.

KIT plays key roles in driving the melanocyte migration from the neural crest along the dorsolateral pathway to colonize the final destination in the skin[17]. Mutations at the *KIT* gene is associated with the Dominant White coat colour of several important commercial breeds, like Large White and Landrace. The Dominant White coat colour is determined by the duplication of about 450-kb region encompassing the entire KIT gene (copy number variation, CNV) and by the presence of a splice mutation in intron 17 in one of the duplicated copies, that causes the skipping of exon 17[3]. *KIT* allele with two normal *KIT* copies has been considered to cause the presence of pigmented regions (patches) in white pigs[3]. In addition, a hypothesis has been proposed that the *KIT* allele carries a single copy of a mutated *KIT* gene (with splice mutation) that should be lethal if homozygous [3, 18]. These perspectives have not yet been validated by functional studies in the past decades. Our functional study confirmed that homozygous of the splice mutation in *KIT* gene is lethal, as *KIT ^D17/D17^* mouse could not be obtained. However, whether the lethal is due to anemia in embryonic stage as previously found in *KIT* defect mouse model need to be further validated. We also found that duplication of the *KIT* gene may not contribute to the patch phenotype found in pigs, as both *KIT ^Dup/+^* and *KIT ^Dup/Dup^* mice did not present the patch coat like that on Pietrain pig, but a coat colour basically indistinguishable from the *KIT^+/+^* control mice (Fig.1C & S1 Fig). Previous study has proposed that increased KIT expression from pig *I^P^* may affect ligand availability, which in turn disturbs the migration of melanocyte precursors, resulting in the patch phenotype [2]. This dosage effect may not be true, as our results showed that increased expression level of KIT protein from additional copy of *KIT* gene (*KIT* CDS in our mouse model) seems to have minimal effect on signaling function of KIT protein receptor on PI3K and MAPK pathways (Fig. 3A & 3C), and thus no obvious effect on melanoblast migration, melanocyte and follicle development, and melanin synthesis. In different to KIT duplication presented in *I^P^* allele in the pig, in our KIT duplication mouse model, only *KIT* CDS was inserted, the large fragment of regulatory regions was not included. The duplicated copy in *I^P^* allele may lack some regulatory elements located more than 150 kb upstream of *KIT* gene body[4], this regulatory mutation may lead to dysregulated expression of one or both copies of KIT, and thus contribute to the patch phenotype. Mutations in other genes responsible for pigmentation in pigs may be associated with the patch phenotype could not be ruled out.

The exon 17 of *KIT* gene encodes the 790-831 amino acids of KIT protein receptor, a highly conserved region of tyrosine kinase domain, which contains Tyr 823 residue that that is conserved in almost all tyrosine kinases, which is phosphorylated during KIT activation and is thought to act to stabilize the stability of KIT tyrosine kinase activity [8]. The splice mutation leading to the lacking of this region is previous considered to be responsible for the impaired KIT signal transduction, and thus the severe defect in the migration and survival of melanocyte precursors. Our IHC analysis of the skin tissue confirmed that the splice mutation can impair KIT signal transduction on PI3K and MAPK pathways (Fig. 4C). PI3K pathway regulates cell growth, proliferation, differentiation and survival [24], and MAPK regulates cell proliferation and apoptosis [25]. In addition, the MAPK pathway is also responsible for phosphorylating and activating MITF, which in turn activates the transcription of mRNAs of several important proteins involved in melanin synthesis, such as Tyrosinase, TRP and TRP2 [19]. The impaired melanoblast migration during embryonic stage (Fig. 4A), and reduced number of melanocyte and melanin accumulation in hair follicle (Fig. 3A) in *KIT ^D17/+^* mice, could be attributed to the impaired PI3K and MAPK signaling induced by the splice mutation.

Previous study proposed an evolutionary scenario whereby *KIT* duplication occurred first and resulted in a white-spotted phenotype, and the splice mutation occurred subsequently and resulted in a completely white phenotype. The presence of one normal *KIT* copy in *I* ensures that white pigs have a sufficient amount of KIT signaling to avoid severe pleiotropic effects on hematopoiesis and germ-cell development [5]. Thus the *KIT* duplication mutation seems to exhibit a rescue function to the splice mutation. Therefore, at first, we expected that the *KIT* duplication could restore the splice mutation in *KIT ^Dup/D17^* mice. However, combination of these two mutations lead to more severely impaired signaling on PI3K and MAPK pathways (Fig.5E & 5F), melanblast migration (Fig. 5H), melanocyte number reduction and melanin accumulation (Fig.5 H), resulting in nearly completely white coat colour (Fig. 5B). Thus the *KIT* duplication mutation does not seems to play a rescue role to the splice mutation. The underlying mechanism of the interaction between these two mutations could be very complicated. As the activation of intrinsic kinase activity of KIT receptor dependents on the binding of SCF ligand to form homodimer. Thus we speculated that improved expression of normal form of KIT protein in *KIT ^Dup/D17^* mice may increase the chance of formation of KIT/KIT ^D17^ dimer upon the binding of SCF as compared with that in *KIT ^D17/+^* mice. Given that the amount of SCF ligand is limited, more KIT/KIT ^D17^ dimer presented on the cell surface of melanoblast may significantly reduce the activation of subsequent PI3K and MAPK signaling pathways, resulting in more severely impaired melanoblast migration, and a more pronounced phenotype change in coat colour (Fig. 6).

Through observation of the light and electron micrograph of a section of skin, a previous concluded that melanocytes and their precursors were absent in the hair bulb of the dominant white (*I*) pigs, and the dominant white color in the pig is due to a defect in the development of melanocytes [20]. However, we found that the combination of *KIT* duplication mutation and splice mutation did not completely block the melanoblast migration (Fig. 5H), and few melanocytes or their precursors can be detected in the skin hair follicles of *KIT ^Dup/D17^* mice through immunostaining of marker protein of melanocytes (Fig. 5C). This was confirmed by q-PCR and Western blot analysis of additional melanocytes marker proteins in the skin of *KIT ^Dup/D17^* mice (Fig. 5D). In addition, in our unpublished experiments, expression of several melanocyte marker proteins was detected by q-PCR and Western blot in skin tissue of Large White pigs, implying the exist of melanocyte or its precursors in dominant white pigs. These results indicate that the combination of KIT duplication mutation and splice mutation although impair the development of melanocyte severely, but still few melanocyte precursors can migrate to destination.

In coclusion, our study provides a further insight into the underlying genetic mechanisms of porcine dominant white coat colour.

## Materials and Methods

### Establishment of mouse models

All mouse models are established on the C57BL/6 background by Model Animal Research Center of Nanjing University (China) as described in previous report [21], with minor modifications. Briefly, C57BL/6 mice were kept under a 12/12 h light/dark cycle. To produce zygotes for pronuclear injection, female mice were injected with 5 IU pregnant mare’s serum gonadotropin (PMSG), and 46–48 h later injected with 5 IU hCG to induce ovulation 10-12 h later. Following the hCG injection, put the females together with male mice in single cages overnight. Fertilized oocytes were isolated from the oviducts for pronuclear injection. To generate *KIT ^Dup/+^* and *KIT ^GtoA/+^*mouse model, Cas9 mRNA, sgRNA and the according donor plasmid (Fig. 1B) were injected into the pronuclei of zygotes. To generate *KIT ^D17/+^*mouse model, Cas9 mRNA and a pair of sgRNA (Fig. 1B) were injected. All sgRNA sequences are listed in S2 table. Injected zygotes were transferred into the oviducts of surrogate recipient female mice to deliver genome-edited pups. *KIT ^Dup/D17^* mice were obtained by mating *KIT ^D17/+^*females with *KIT ^Dup/+^* males because *KIT ^D17/+^*males are infertile.

All procedures were performed in strict accordance with the recommendations of the Guide for the Care and Use of Laboratory Animals of the National Institutes of Health. The protocol was approved by the Institutional Animal Care and Use Committee (IACUC), Sun Yat-sen University (Approval Number: IACUC-DD-16-0901).

### Mouse genotyping

Mice genotypes are identified by PCR. The tail of 1week old mice are cut off to extract DNA by using a tissue DNA extraction kit (OMEGA). Primers used in PCR are summarized in S2 table.

Mice skin RNAs were prepared using TRIzol (Invitrogen) extraction followed by DNase (Ambion) treatment, and reverse transcription was carried out using the instructions of Reverse Transcription System (Promega).

Primers for mouse genotype identification please refer to S2 table. Polymerase chain reaction (PCR) was carried out using the GeneStar^TM^ PCR Mix system. Each PCR reaction mix contained 1× GeneStar^TM^ PCR Mix buffer, 1.0µM of each primer and about 100 ng DNA template. The procedure in the thermal cycling was an initial 5 min hold at 95 °C, followed by 35 cycles of 30 sec at 95°C, 30 sec at 60°C, and 30 sec at 72°C, and finishing with 10 min incubation at 72°C.

In order to determine the mutant sequences, the PCR products were recovered by OMEGA DNA Gel Recovery Kit, cloned into pMD-18T vector (TAKARA) and transformed into DH5α competent cells. Plasmids then were purified from *E. coli* cells for Sanger sequencing.

### Isolation and culture of peritoneal cell derived mast cells

Isolation and culture of peritoneal cell derived mast cells was performed as previous described protocol [22] with minor modifications. 5 ml of PBS and 2 ml of air was injected into peritoneal cavity of 14 weeks old mice. Then injected mice were carefully shaken in the palm for 5 mins. Subsequently, the cell-containing fluid of the peritoneal cavity is gently collected in a plastic Pasteur pipette. After centrifugation, cells were resuspended, and cultured in DMEM medium supplemented with serum, cytokines IL-3 (CP39; novoprotein) and SCF (C775; novoprotein). After 10 days culture, CD117 and FceR1 makers were used to determine whether cultured cells are mast cell by flow cytometry analysis using the Beckman Coulter Gallios™ Flow Cytometer. For surface staining, cells were stained with APC anti-Mouse CD117 (17-1171; affymetrix) and FITC anti-mouse Fc epsilon receptor I alpha (FceR1) (11-5898; affymetrix) at room temperature for 30 minutes, then washed with PBS and then re-suspended in PBS.

### Histological and immunohistochemical analysis of tissue sections

Skin tissues and embryos were fixed overnight in 10% (w/v) paraformaldehyde with 0.02 MPBS (pH 7.2) at 4 °C, processed and mounted in paraffin, then serially cut into 5μm-thick sections by Rotary Microtome (MICROM). Histological sections were stained with hematoxylin and eosin (H&E), observed and photographed under a fluorescent microscopy (Zeiss). For immunohistochemistry experiments, sections were treated with 3% H_2_O_2_ to quench endogenous peroxidase activity, then treated with 5% bovine serum albumin to block nonspecific protein binding sites. Sections were incubated with primary antibody at 4 ◦C overnight, and then stained by using anti-Rabbit HRP-DAB Cell and tissue staining kit (R&D, CTS005). Detection was followed by TSA plus Fliorescein (Perkinelemer, NEL741001KT). All antibodies including KIT (ab47587; abcam), Phospho-KIT (Try719) (#3391; Cell Signaling), Green Fluorescent Protein (AB3080P; merk), DCT (ab74073; abcam), Erk1/2 (#4695; Cell Signaling), Akt (#9272; Cell Signaling) Phospho-Akt (#4060; Cell Signaling) and Phospho-Erk1/2 (#8544; Cell Signaling) were diluting in 1:200 with PBS.

### qPCR

For all gene expression level detection, total RNAs were prepared using TRIzol (Invitrogen) extraction followed by DNase (Ambion) treatment, and reverse transcription was carried out following the instructions of Reverse Transcription System (Promega). The resulting total cDNAs were analyzed quantitatively using FastStart Universal SYBR Green Master kit (Roche) with primers for *KIT, ERK, AKT, PLCG* and *DCT*. Expression profiles were tested in triplicate on at least two mice on an LC480 instrument (Roche). Data were analyzed using the comparative Ct (ΔΔCt) method and one-tail, unpaired student T test (significance cutoff p<0.01). Gene expression levels were normalized to the housekeeping gene glyceraldehyde 3-phosphate dehydrogenase (GAPDH).

### Western blot analysis

Proteins were extracted using Lysis Buffer (Key GEN), and proteins concentration was determined by using PierceTM BCA Protein Assay Kit (Thermo). 300 ng protein was subjected to 10% SDS gel and electrotransferred onto PVDF membrane (Roche). After blocking for 1h with 3% BSA in PBS, the membrane was incubated with primary antibodies at 4 ◦C overnight.

The rabbit anti-KIT antibody (ab47587; abcam), rabbit anti-Phospho-KIT (Try719) antibody (#3391; Cell Signaling), rabbit anti-Green Fluorescent Protein antibody (AB3080P; merk), rabbit anti-DCT antibody (ab74073; abcam), rabbit anti-Erk1/2 antibody (#4695; Cell Signaling) and rabbit anti-Akt antibody (#9272; Cell Signaling) were diluting in 1:1000, rabbit anti-Phospho-Akt antibody (#4060; Cell Signaling) and rabbit anti-Phospho-Erk1/2 antibody (#8544; Cell Signaling) were diluting in 1:2000, rabbit anti-GAPDH antibody (AP0063; biogot) was diluting in 1:5000 with PBS. Following by 10 min three times washing with TBST, the membrane was then incubated with 1:5000 goat anti-rabbit secondary antibodies (Abcam, ab6721) for 1h at room temperature. Protein bands were visualised using Kodak image station 4000MM/Pro (Kodak), according to the manufacturer’s instructions, and exposed to FD bio-Dura ECL (FD, FD8020). Protein levels were standardized by comparison with GAPDH.

### Statistical analysis

All data were analyzed by using EXCEL (version 2016). The data were expressed as the means ± SEM. Only values with p < 0.05 were accepted as significance.

## Acknowledgement

This work was jointly supported by National Transgenic Major Program (2016ZX08006003-006), National Key R&D Program of China (2018YFD0501200), and Key R&D Program of Guangdong Province (2018B020203003).

## Supporting information

**S1 Fig. *KIT ^GtoA/+^, KIT ^D17/+^* and *KIT ^Dup/Dup^* mice phenotype.** (A) White spots appeared on the abdomen of *KIT ^GtoA/+^ mice as compared with KIT^+/+^. KIT ^D17/+^* mice presented a piebald coat colour in head and trunk, a vertical white stripe on the forehead, a half loop of white hair on the shoulder blade area, and dominant white at entire abdominal region. There was no difference between the coat colour of *KIT^+/+^* and *KIT ^Dup/Dup^* mice at 14 W old. (B) Schematic diagram of primers designed for identification of *KIT ^D17/+^* and *KIT ^Dup/Dup^* mice identification. (C) PCR identification of the *KIT ^D17/+^* and *KIT ^Dup/Dup^* _mice._

**S2 Fig. Identification of mouse peritoneal mast cells through flow cytometry analysis.** KIT (stained by anti-Mouse CD117 APC) and FcεRI (stained by anti-mouse Fc epsilon receptor I alpha) were used as markers of mast cell.

**S3 Fig. Using KIT as marker to detect melanoblast migration in *KIT ^+/+^* and *KIT ^Dup/+^* mice embryo.** KIT is used as marker to detect melanoblast migration in *KIT ^+/+^* and *KIT ^Dup/+^* mice embryo (E14.5).

**S4 Fig. Coat colour changing of *KIT ^Dup/D17^* mice during growth up.** With the increase of age, the gray hairs of the eyelids and hips gradually disappeared.

**S1 Table. Homologous of splice mutation of KIT gene could be lethal.** Oocytes from superovulated *KIT ^D17/+^* females were *in vitro* fertilized with sperms collected from 5 *KIT ^D17/+^* males, and transferred to 10 surrogate females to generate offspring. No *KIT ^D17/D17^* pups were born but 13 *KIT ^+/+^* and 24 *KIT ^D17/+^* pups were obtained.

**S2 Table. Oligos and primers used in this study.**

## References

1. Wiseman J. A history of the British pig: Duckworth; 1986.

2. Marklund S, Kijas J, Rodriguez-Martinez H, Rönnstrand L, Funa K, Moller M, et al. Molecular basis for the dominant white phenotype in the domestic pig. Genome research. 1998;8(8):826–33.

3. Pielberg G, Olsson C, Syvänen A-C, Andersson L. Unexpectedly high allelic diversity at the KIT locus causing dominant white color in the domestic pig. Genetics. 2002;160(1):305–11.

4. Giuffra E, Törnsten A, Marklund S, Bongcam-Rudloff E, Chardon P, Kijas JM, et al. A large duplication associated with dominant white color in pigs originated by homologous recombination between LINE elements flanking KIT. Mammalian Genome. 2002;13(10):569–77.

5. Andersson L, editor Studying phenotypic evolution in domestic animals: a walk in the footsteps of Charles Darwin. Cold Spring Harbor symposia on quantitative biology; 2010: Cold Spring Harbor Laboratory Press.

6. Rubin C-J, Megens H-J, Barrio AM, Maqbool K, Sayyab S, Schwochow D, et al. Strong signatures of selection in the domestic pig genome. Proceedings of the National Academy of Sciences. 2012;109(48):19529–36.

7. Lennartsson J, Rönnstrand L. Stem cell factor receptor/c-Kit: from basic science to clinical implications. Physiological reviews. 2012;92(4):1619–49.

8. Roskoski R. Structure and regulation of Kit protein-tyrosine kinase—the stem cell factor receptor. Biochemical and biophysical research communications. 2005;338(3):1307–15.

9. Imokawa G. Autocrine and paracrine regulation of melanocytes in human skin and in pigmentary disorders. Pigment cell research. 2004;17(2):96–110.

10. Wehrle-Haller B, Weston JA. Receptor tyrosine kinase-dependent neural crest migration in response to differentially localized growth factors. BioEssays. 1997;19(4):337–45.

11. Rubin BP, Antonescu CR, Scott-Browne JP, Comstock ML, Gu Y, Tanas MR, et al. A knock-in mouse model of gastrointestinal stromal tumor harboring kit K641E. Cancer research. 2005;65(15):6631–9.

12. Moniruzzaman M, Sakamaki K, Akazawa Y, Miyano T. Oocyte growth and follicular development in KIT-deficient Fas-knockout mice. Reproduction. 2007;133(1):117–25. doi: 10.1530/REP-06-0161. PubMed PMID: 17244738.

13. Bernex F, De Sepulveda P, Kress C, Elbaz C, Delouis C, Panthier J-J. Spatial and temporal patterns of c-kit-expressing cells in WlacZ/+ and WlacZ/WlacZ mouse embryos. Development. 1996;122(10):3023–33.

14. Slominski A, Paus R, Plonka P, Chakraborty A, Maurer M, Pruski D, et al. Melanogenesis during the anagen-catagen-telogen transformation of the murine hair cycle. Journal of investigative dermatology. 1994;102(6):862–9.

15. Hou C, Miao Y, Wang X, Chen C, Lin B, Hu Z. Expression of matrix metalloproteinases and tissue inhibitor of matrix metalloproteinases in the hair cycle. Experimental and therapeutic medicine. 2016;12(1):231–7.

16. Botchkareva NV, Botchkarev VA, Gilchrest BA. Fate of melanocytes during development of the hair follicle pigmentary unit. The journal of investigative dermatology Symposium proceedings. 2003;8(1):76–9. doi: 10.1046/j.1523-1747.2003.12176.x. PubMed PMID: 12894999.

17. Besmer P, Manova K, Duttlinger R, Huang EJ, Packer A, Gyssler C, et al. The kit-ligand (steel factor) and its receptor c-kit/W: pleiotropic roles in gametogenesis and melanogenesis. Dev Suppl. 1993:125–37. PubMed PMID: 7519481.

18. Johansson A, Pielberg G, Andersson L, Edfors-Lilja I. Polymorphism at the porcine Dominant white/KIT locus influence coat colour and peripheral blood cell measures. Animal genetics. 2005;36(4):288–96. doi: 10.1111/j.1365-2052.2005.01320.x. PubMed PMID: 16026338.

19. Lin JY, Fisher DE. Melanocyte biology and skin pigmentation. Nature. 2007;445(7130):843.

20. Moller MJ, Chaudhary R, Hellmen E, Höyheim B, Chowdhary B, Andersson L. Pigs with the dominant white coat color phenotype carry a duplication of the KIT gene encoding the mast/stem cell growth factor receptor. Mammalian Genome. 1996;7(11):822–30.

21. Tao FF, Yang YF, Wang H, Sun XJ, Luo J, Zhu X, et al. Th1-type epitopes-based cocktail PDDV attenuates hepatic fibrosis in C57BL/6 mice with chronic Schistosoma japonicum infection. Vaccine. 2009;27(31):4110–7. doi: 10.1016/j.vaccine.2009.04.073. PubMed PMID: 19410625.

22. Meurer SK, Ness M, Weiskirchen S, Kim P, Tag CG, Kauffmann M, et al. Isolation of Mature (Peritoneum-Derived) Mast Cells and Immature (Bone Marrow-Derived) Mast Cell Precursors from Mice. PLoS One. 2016;11(6):e0158104. doi: 10.1371/journal.pone.0158104. PubMed PMID: 27337047; PubMed Central PMCID: PMCPMC4918956.

